# Cropbox: a declarative crop modeling framework

**DOI:** 10.1101/2022.10.10.511649

**Authors:** Kyungdahm Yun, Soo-Hyung Kim

## Abstract

Crop models mirror our knowledge on crops *in silico*. Therefore, crop modeling inherits common issues of software engineering and often suffers from technical debts. We introduce a new crop modeling framework: Cropbox as a declarative domain-specific language. Recognizing that a crop model is fundamentally an integrated network of generalized state variables, we developed the framework to encapsulate abstract primitives for representing variables, systems, and functions that are essential to crop modeling workflows. With a constrained syntax, high-level model specifications are automatically translated into low-level host code written in Julia programming language. This allows complex crop models to become more accessible and transparent for modelers to build and use. We highlight key capabilities of the Cropbox framework through specific case studies featuring a coupled leaf gas-exchange model and a process-based crop simulation model. We also illustrate potential extensions of the framework to support functional-structural plant modeling (FSPM) using a 3D root architectural model as an example.

## 1 Introduction

### 1.1 Crop Modeling

Crop modeling is inherently a software engineering process that the model as an outcome is subject to technical implications posed by the platforms and tools used for developing the model. The history of crop modeling has shown multiple approaches influenced by development paradigms available at the time.

Early crop models from Wageningen University were originally developed on CSMP (Continuous System Modeling Program) language which was a simulation language designed for continuous time domain where problems are represented by differential equations. CSMP was based on the idea of clear separation between groups of variables involved in numerical integration [Brennan and Silberberg, 1968, Caskie and Mason, 1972]. Rate variables represent changes of certain quantities during a small time step. State variables hold quantities at each time point integrated by the rate variables as integrands.

Once the language became obsolete with the arrival of new computer platforms in 1980s, an attempt to replace CSMP was conceived to continue the development of crop models and FSE/FST came out as a result. FSE (Fortran Simulation Environment) was a framework equipped with domain-specific simulation features similar to CSMP on top of FORTRAN which was a general programming language widely used at the time [van Kraalingen and de Vries, 1990, van Kraalingen, 1991, 1995]. FST (Fortran Simulation Translator) was a companion tool providing a transition layer for existing models written in CSMP syntax [Rappoldt and van Kraalingen, 1996, van Kraalingen et al., 2003]. Variables were still declared in separate sections in the code to distinguish rate and state variables. Order of calculation within the section was automatically determined by sorting out the dependency of the variables involved. In addition to the model specification mainly describing a set of differential equations, modelers also need means of running the models and evaluating the outcomes. For a complete simulation workflow, FSE/FST provided model driving statements in order to support setting up parameters, including time step and output format, and producing graphical plots. FSE/FST being a domain-specific simulation language implemented on Fortran since then became widely adopted for developing varieties of crop models. A major revision of DSSAT (Decision Support System for Agrotechnology Transfer) was once driven by a transition to FSE/FST framework and the internal model structure established at the time largely remains to date [Jones et al., 2003]. With the advent of recent technological shifts, FSE has also inspired new simulation frameworks implemented in different programming languages. PCSE (Python Crop Simulation Environment) was an attempt to reimplement existing crop models written for FSE in a dynamically typed language Python which could be more suitable for interactive simulation [de Wit et al., 2019]. WISS (Wageningen Integrated Systems Simulator) was a recently announced simulation framework developed in Java language with a similar objective in mind [van Kraalingen et al., 2020].

### 1.2 System Dynamics

System dynamics is an approach to understand emergent behavior of complex systems by linking simple primitives like stock and flow [Forrester, 1961]. Mathematically, system dynamics model consists of a set of differential equations similarly to continuous system simulation languages. Stock, or originally called level, represents a quantity accumulated or integrated over time. Flow represents a change of stock per time which is essentially the same concept as rate variable above. Incoming flow and outgoing flow are explicitly identified as separate entities to clearly show interactions between variables. System dynamics model can be visualized by a diagram with own set of unique symbols for representing stock and flow. Quantities used in the model are often accompanied by proper units of measurements which are useful for validating model equations. The basic concept of system dynamics was established throughout revisions of DYNAMO (DYNAmic MOdels) language in 1960s [Forrester, 1961] and later evolved by other platforms. STELLA (Systems Thinking, Experimental Learning Laboratory with Animation) started as an implementation of DYNAMO for the Macintosh computers in 1980s and provided a graphical user interface supporting model development directly in the form of stock–flow diagrams [Richmond, 1985]. Many ecological models were implemented on STELLA [Costanza et al., 1998, Costanza and Gottlieb, 1998, Costanza and Voinov, 2001]. Simile is a visual modeling environment primarily based on the concept of system dynamics with additional features of object-based paradigm suited to spatial and individual-based modeling [Muetzelfeldt and Massheder, 2003]. Boosd (Bergen Object-Oriented System Dynamics) is a textual language for system dynamics with an object-oriented extension [Powers, 2011].

### 1.3 Object-Oriented Programming

Object-oriented programming (OOP) was adopted by some crop models for managing complexity of the model [Silvert, 1993, van Evert and Campbell, 1994, Acock and Reddy, 1997, Sequeira et al., 1997, Lemmon and Chuk, 1997]. Earlier imperative programming approaches did not provide a clear boundary between a group of variables and functions that work on them. OOP helped organize a complex model structure via encapsulation of data and the associated code in the structure called object. For instance, an object in a crop model could implement a certain physiological process or could represent a biological structure. As a new child object may be composed of existing objects and share the same interface as the parent, some objects can be reused in other models. In other words, crop models became modular and the models were developed by integration of multiple components which could be easily replaced by another components when necessary [Reynolds and Acock, 1997, Papajorgji et al., 2004, Argent, 2004, Donatelli et al., 2006, Holzworth et al., 2010, Adam et al., 2012]. Examples of modular modeling framework include 2DSOIL [Timlin et al., 1996], DSSAT [Jones et al., 2001], ModCom [Hillyer et al., 2003], OpenAlea [Pradal et al., 2008], OMS (Object Modeling System) [David et al., 2013], Universal Simulator [Holst, 2013], VLE (Virtual Lab Environment) for RECORD used by STICS [Bergez et al., 2013, 2014], and Plant Modeling Framework (PMF) for APSIM [Brown et al., 2014, Holzworth et al., 2014, 2018]. Functional-structural plant modeling (FSPM) frameworks, such as FSPM-P [Henke et al., 2016], Helios [Bailey, 2019], and CPlantBox [Zhou et al., 2020], also adopted OOP for coordinating a large number of homogeneous structural components. At a larger scale, demand for integrating multiple individual models across different languages and platforms arose recently. Yggdrasil provided a template for describing data exchange between standalone models implemented in different languages [Marshall-Colon et al., 2017, Lang, 2019]. Crop2ML allowed the reuse of model components developed in a domain-specific language called CyML by automatically generating code for target model platforms [Midingoyi et al., 2020, 2021].

### 1.4 Declarative Modeling

Early simulation languages, like CSMP and FSE/FST, and most implementations of system dynamics mentioned above embraced a declarative modeling approach tailored for each domain. Soon the increased complexity of modeling triggered a paradigm shift and led to the adoption of more general programming languages, seeking solutions from a software engineering perspective. For instance, component-based model development supported by object-oriented programming (OOP) gained momentum for building large-scale models. However, the reusability of the model was still limited, especially when model components had overlapping internal states and imperative control flows. Declarative modeling is an alternative approach representing models in a high-level abstraction focusing on relations (“what”) rather than procedures (“how”) which are implementation details that can be hidden away [Muetzelfeldt, 2004]. Many domain-specific languages therefore naturally followed the style of declarative modeling [Athanasiadis and Villa, 2013] as evidenced by MOSES (MOdelling and SImulation of Ecological Systems) [Wenzel, 1992], SIGMA (Scientists’ Intelligent Graphical Modeling Assistant) [M Keller and L Dungan, 1999], ECOBAS [Benz et al., 2001], Integrating Modelling Architecture (IMA) [Villa, 2001], and ECHSE [Kneis, 2015].

### 1.5 Crop Modeling Framework

The role of crop modeling framework from a software development perspective has been discussed multiple times in the past. An early review suggested that a modeling framework should provide common services essential for implementing a model such as state variable integration and event handling [van Ittersum et al., 2003]. Other common features included a graphical user interface and a standard format for handling data input and output. By separating core simulation logic from other supporting code, models would become more independent and modular for better reuse. A later review revisited the state of crop modeling development highlighting that many long-standing problems were still present after a decade of progress in the field [Holzworth et al., 2015]. Crop models still relied on legacy code written in procedural languages which might not represent the most effective paradigm for implementing scientific simulation models. A standard model interface and internal structure with smaller units of computation aggregated to form a large model were suggested with an emphasis on documentation and testing. A recent review led to a similar conclusion that crop modeling should follow best practices established in software engineering such as adopting domain-specific structures and functions and clear separation of code between model equation and user interface [Janssen et al., 2017].

In this paper, we introduce a new crop modeling framework named Crop-box that allows models to be expressed in a domain-specific language akin to mathematical model equations where each variable is declared with an explicitly stated intention and relation to other variables. Model developers can write a model specification in a declarative form with no direct need for control flow management as in procedural languages. The Cropbox framework then interprets this high-level model specification and automatically generates lower-level code in a host language Julia after analyzing the dependency graph between variables. Model developers and users can use built-in features including, but not limited to, running simulation of the model, configuration management, evaluation with common metrics, calibration of parameters, visualization of output results, and manipulation of interactive plots. A core concept and architecture of the framework will be discussed with a few applications taking advantage of different aspects of the framework (Figure 1).

**Figure 1:**
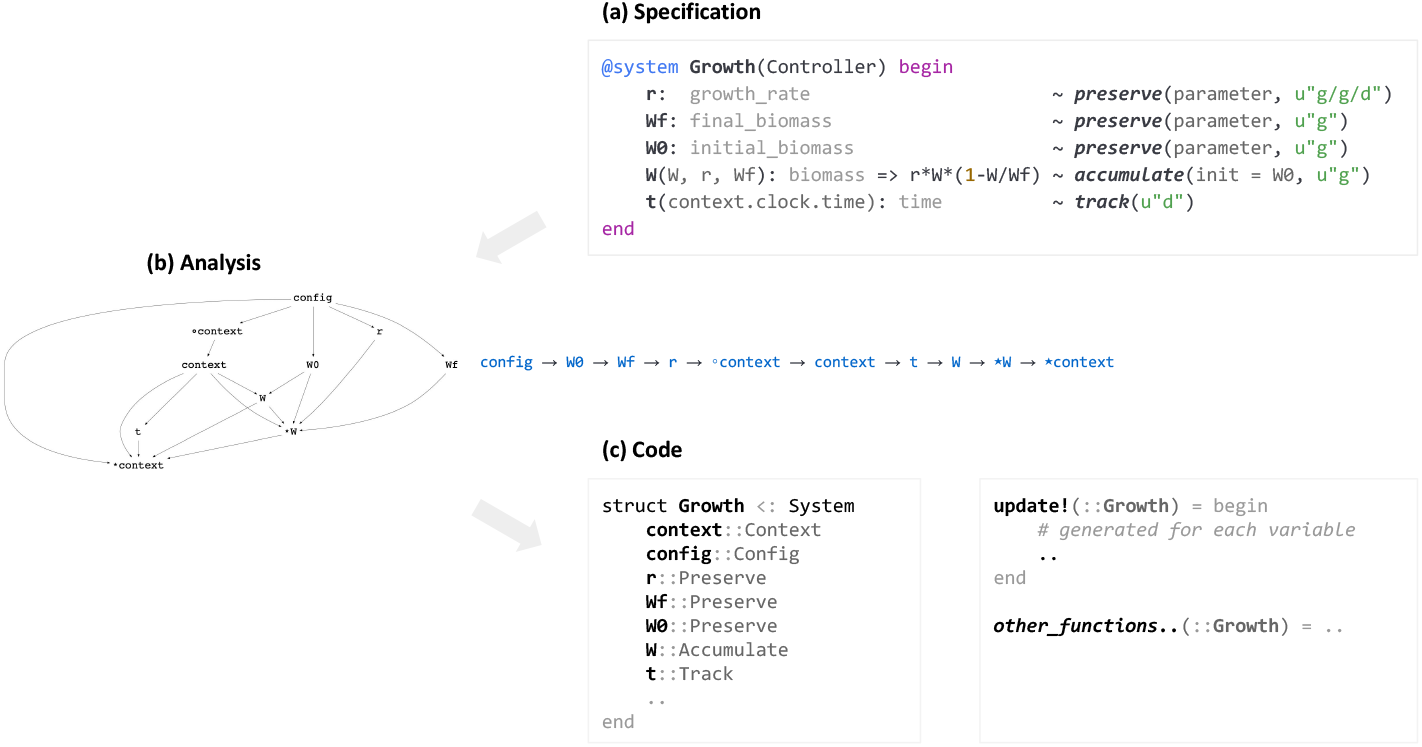
Concept of Cropbox modeling framework. The framework takes a model specification in declarative domain-specific language and emits host code written in Julia programming language after internal dependency analysis for variables and systems. Model developers then interact with automatically generated model code via several functions provided for simulation, evaluation, calibration, and visualization workflow.

## 2 Architecture

### 2.1 Variable

A model in the Cropbox framework is represented by relations between variables. Variable is a unit element of modeling that denotes a value determined by a specific operation relying on other variables. The framework provides more than dozens of predefined operations (Section 2.1.2) from a simple constant value assignment to a mathematical equation to a specialized computational code. Operation is internally called *state* in terms of generalizing and expanding the notion of *state variable* as often mentioned in mathematical modeling. State variable, in an original strict term, captures a state of integration upon a dynamic variable with regard to the time variable. In our broad terms, we assume more types of states exist to conveniently describe how a model should work in multiple aspects. Associating a single type of predefined operation with each variable makes model developers free from technical details of how an operation is implemented under the hood and instead focus on what kind of value a variable should keep to describe a desired aspect of the model. Such a clear separation between high-level model specification and low-level computer code is a core idea of the Cropbox modeling framework. Model developers are encouraged to think around variables as appeared in mathematical equations which they want to translate into a computer model.

#### 2.1.1 Syntax

A state variable in Cropbox is declared in a minimal form like `name ∼ state`. Most variables may have additional descriptions inside the body as in `name => body ∼ state`. Depending on variable types, the body may include a single value, a set of equations to calculate a new value, or a block of regular function calls incurring side effects. When references to other variables are needed, those dependent variables are listed as *arguments* appearing next to the variable name as in `name(args…) => body ∼ state`.

Note that, while the form of `name(args…) => body` may look like a function and there is indeed a special occurrence when this form gets translated into a function-like behavior (*i*.*e*. `call` variable), it normally just indicates a dependence relationship that the value of variable named `name` should be determined by other variables listed in `args`.

Similarly, state-specific options, called *tags*, may optionally appear as arguments of the state name as in the form of `name => body ∼ state(tags…)`. Commonly used tags include unit specifier (`u”..”`), parameter indicator (`parameter`), and clamping range (`min`, `max`). Variable may have an *alias* which is a longer description complementary to the original name which is encouraged to be kept short for the sake of convenience when referred by other variables. An internal data type can be also specified when default floating point type is not enough, for example, when a strict integer type is needed for counting. Note that specifying *state* is not mandatory when declaring variables and the use of plain variables are also allowed. Such non-state variables are often used for referencing another system which is a container of variables.

#### 2.1.2 States

There are currently 19 types of states implemented in the framework (Table 1). Some variables describing essential building blocks of the model are more common while others are more specialized to handle unique but complex tasks. Here we have a brief overview of commonly used state variables.

**Table 1:** List of state types currently implemented in the framework.

| State | Description |
| --- | --- |
| *Instant derivation* |  |
| **preserve** | keeps an initially assigned value with no further updates; constants, parameters |
| **track** | evaluates expression and assigns a new value for each time step |
| **flag** | checks a conditional logic; similar to **track** with boolean type, but composition is allowed |
| **remember** | keeps tracking the variable until a certain condition is met; like **track** switching to **preserve** |
| *Cumulative update* |  |
| **accumulate** | emulates integration of a rate variable over time; essentially Euler method |
| **capture** | calculates the difference of **accumulate** between time steps |
| **integrate** | calculates an integral over a non-time variable using Gaussian method |
| **advance** | updates an internal time-keeping variable |
| *Data source* |  |
| **provide** | provides a table-like multi-column time-series data; *i.e.* weather data |
| **drive** | fetches the current value from a time-series; often used with **provide**; *i.e.* air temperature |
| **tabulate** | makes a two dimensional table with named keys; *i.e.* partitioning table |
| **interpolate** | makes a curve function interpolated with discrete values; *i.e.* soil characteristic curve |
| *Equation solving* |  |
| **solve** | solves a polynomial equation symbolically; *i.e.* quadratic equation for coupling photosynthesis |
| **bisect** | solves a nonlinear equation using bisection method; *i.e.* energy balance equation |
| *Dynamic structure* |  |
| **produce** | attaches a new instance of dynamically generated system; *i.e.* root structure growth |
| *Language extension* |  |
| **hold** | marks a placeholder for the variable shared between mixins |
| **wrap** | allows passing a reference to the state variable object, not a dereferenced value |
| **call** | defines a partial function accepting user-defined arguments, while bound to other variables |
| **bring** | duplicates variables declaration from another system into the current system |

`preserve` is used when a value of the variable should be kept constant with no further modification after the initial assignment. Hence it is often used for declaring parameter variables where the initial value can be set via a configuration object supplied with the onset of the simulation. Non-parameter constant variables may also appear depending on model specification, for example, to freeze a value from a certain time point and lower computational cost since no update is required for each time step in contrast to other types of variables like `track`.

~~~
“Relative growth rate of plant biomass.”
r: growth_rate => 0.05 ∼ preserve(parameter, u”d^-1”)
~~~

`track` is for tracking a result of computation as we would do in an update loop in conventional model code. For each time step, the model equation coded in the variable declaration body is evaluated and saved for use by other variables. Note that the assignment of value can happen only once per time step as determined by the framework and no manual assignment at an arbitrary point of computation is allowed as with procedural programming. This *declarative* nature of the framework is to prevent any logical errors resulting from premature or incorrectly ordered variable assignments which can happen frequently in model development. For example, two variables declared to track each other with a circular reference would be easily caught by the framework during the model analysis phase before code generation. Such kind of logical errors would be treated as valid in procedural programming languages, only leading to bugs that are hard to detect. Some of these subtle errors may never be discovered even after the model is deployed and while used in production.

~~~
“Specific leaf area, which is the inverse of LMA.”
SLA(area, mass): specific_leaf_area => **begin**
  area / mass
**end** ∼ track(u”cm^2/g”)
~~~

`flag` is basically equivalent to `track` exclusive for boolean values representing either true or false, which are often used for representing discrete events associated with agricultural management such as sowing and harvesting. In addition to tracking a conditional value for each time step, `flag` variable can be easily composed with other flag variables to build a new conditional expression. For example, an expression like `appeared & !removed` indicates a condition when organs like a flower bud, fruits, and scape have appeared but have not been removed or harvested yet. Triggering alternative computational pathways depending on the current condition on the fly is common in crop models, especially when backed with phenology models accounting for complex developmental stages [Yun et al., 2017].

~~~
“The number of leaves appeared.”
leaf_appeared(i = N_initiated, a = N_appeared) => **begin**
  0 < i <= a
**end** ∼ flag
~~~

`accumulate` represents an integration of differential, or more precisely, difference equations over time. The numerical integration is based on the Euler method with a fixed time step since other adaptive numerical methods relying on varying time steps are not compatible with the discrete nature of certain variables like `produce` where conditions for generating a new system instance should be checked every time step and the checking cannot be easily skipped when a larger adaptive time step is used.

The declaration body of `accumulate` variable contains an equation describing how the accumulation rate is calculated. Any type of variable can become a part of an integrating variable or an associated time variable if the use of units is consistent. As the framework is equipped with native support for unit conversion and validation, the units of the rate variable must conform to the units of the integrated variable. For example, if biomass is accumulated in the unit of g, the corresponding rate variable must be in the unit with a time component such as g d^−1^. The rate is calculated at the end of each time step, but actual accumulation occurs at the beginning of the following time step to ensure a newly updated value is not accidentally leaked to external access at the end of a previous time step. For convenience, a similar variable type named `capture` is also provided to allow tracking of rate values calculated for each time step.

~~~
“Total biomass of the plant.”
W(r, W, Wf): biomass => **begin**
  r * W * (Wf - W) / Wf
**end** ∼ accumulate(u”g”, init = W0)
~~~

`drive` pulls the value from a time-series data source which can be a simple vector or a column from a data frame. Depending on the internal data type of the specified index variable which often coincides with the default time or calendar date variable, new values are pulled from the source every time step. In other words, it declares a *driving variable* hence the name. When the data source is in a data frame, `provide` automatically generates a convenient interface for loading from external files, handling index columns, and automatic unit conversion from column names.

~~~
“Hourly air temperature in the degree Celsius.”
Ta: air_temperature ∼ drive(from = s, by = :T, u”^*?*^C”)
~~~

Aside from basic variable types introduced so far that are more frequently used in general model development, other variables are reserved for abstracting specialized tasks often complex enough to implement from scratch.

For example, `bisect` constructs a loop for iteratively solving a numerical equation between variables. With a lower and upper boundary of the variable to be solved given in the declaration, the framework would automatically generate a proper control flow structure for implementing an iterative solver. When multiple equations are interdependent and declared with more than a single `bisect` variable, dependency analysis can parse out the correct order of iterative solution embedded in a nested control structure. For example, we used `bisect` to solve intercellular CO_2_ level (*C*_*i*_) determined by an empirical relationship between biochemical photosynthesis and stomatal conductance models [Yun et al., 2020]. Additionally, our energy balance equation was solved via another `bisect` variable internally relying on *C*_*i*_ that led to a nested loop generated in the host code.

~~~
“““
CO2 concentration inside the leaf.
The value is determined by coupling,
- CO2 concentration in the air (Ca)
- Net photosynthesis rate (A_net)
- Leaf conductance for CO2 (gvc)
“““
Ci(Ca, Ci, A_net, gvc): intercellular_co2 => **begin**
  Ca - Ci == A_net / gvc
**end** ∼ bisect(min = Cimin, upper = Cimax, u”*μ*bar”)
~~~

`solve` is another type of variable for solving an equation when the target equation is symbolically solvable to produce an analytical solution. Since actual analysis for solving an equation occurs during the code generation phase and only the solution represented in a new fixed form is included in the generated code, the impact on runtime performance should be kept minimal. For example, we used `solve` to calculate leaf surface humidity (*H*_*s*_) in the stomatal conductance model which is a key for coupling with the biochemical photosynthesis model [Yun et al., 2020].

~~~
“““
Photosynthetic electron transport rate.
It controls RuBP-regeneration limited photosynthesis.
“““
J(I2, Jmax, *θ*): electron_transport_rate => **begin**
  a = *θ*
  b = -(I2 + Jmax)
  c = I2 * Jmax
a*J^2 + b*J + c == 0
**end** ∼ solve(lower = 0, upper = Jmax, u”*μ*mol/m^2/s”)
~~~

`produce` supports the instantiation of a new system during runtime to allow dynamic expansion of complex model structure which is an important feature differentiating functional-structural plant models (FSPM) from conventional crop models with a rather static structure [Vos et al., 2010, Soualiou et al., 2021]. We used `produce` to build a 3D root structure growth model for switchgrass (*Panicum virgatum*) based on the root growth algorithm described by CRootBox model [Schnepf et al., 2018].

~~~
“A collection of axial root segments.”
R(R, N, wrap(RT0)): roots => **begin**
  [produce(Root; RT0) **for** i **in** (length(R)+1):N]
**end** ∼ produce::Root[]
~~~

### 2.2 System

System is a unit of model component that contains a collection of variables declared with metadata as described above. The framework guarantees that the most current state of variables is readily accessible by other variables in the same system when those depending variables are being referenced. To ensure a correct propagation of variable states, the system must have a linear order of computation that satisfies all the requirements for dependency imposed by variable declarations. If any inconsistency caused by a cyclic dependency among some variables were found during the dependency analysis phase, code generation would be halted and an error would be thrown. Variables declared in another system can be also accessed if the entire system holding dependent variables has been already updated. In other words, an external system can be declared as a member of the system which becomes a node in the dependency graph. The resolved order of computation then should reflect a valid timing of updates for other systems.

A system can be viewed as a class in object-oriented programming that a definition of a certain system works like a blueprint for imprinting an instance of the system where actual storage for keeping the states of variables are allocated in the computer’s memory. Models declared in Cropbox first need to have underlying systems instantiated before running any simulation.

For complex models, a system may be composed of other existing systems in a sense that the system *has*, or includes, variable declarations simply copied from other systems. It is like multiple blueprints of systems are all combined to form a large single template. For example, a gas-exchange model may be defined as a system named `LeafGasExchange`, which is composed of multiple components such as `C3Photosynthesis`, `StomataBallBerry`, `EnergyBalance` and likes, exports all variables from these components in a merged namespace.

Cropbox adopts *composition* as a primary method for modular model development. Systems that are reused as components for building a new system are called *mixins* since they are mixed in to make a new system while avoiding complicated parent-child relationships [Bracha and Cook, 1990]. If desired, an existing mixin component of the system can be substituted with another mixin at the time of system declaration to implement a modular plugin architecture. It is like a new mixin can be plugged into the place of an old mixin which is replaceable if both mixins share a similar interface. A new system built by composition includes all variables declared in the original systems, but the resulting system itself is not recognized by any type signatures from constituting components. Technically, the new system is not a child, or subtype, of the components and therefore needs to be referenced by its own unique type.

Specific modeling aspects are often better represented by a strict hierarchy. Some systems may share common traits while they are still distinct enough to deserve their own system types. Such systems are often collectively referred to by the common parent type as if they belong to the same type. When such polymorphism is desired, a system can designate an optional base type that is inherited by the new system. Such a relationship expresses that the new system *is* essentially a kind of its parent system in the sense that the type of child system is inherited from the type of parent system. Root structure can be a good example of hierarchical modeling where different types of root segments are inherited from the same `BaseRoot` system sharing declaration of common variables, like length and angle, and each system is declared as a subtype of `BaseRoot`, namely `PrimaryRoot`, `FirstOrderLateralRoot`, and `SecondOrderLateralRoot`, to model a specific behavior of growth for each root type. Without such an explicit declaration of a base type, systems are all inherited by abstract `System` type by default in Cropbox. Inheritance can be useful for designing a special interface around a hierarchy of custom system types. The root structure model would rely on `produce` variable for dynamically generating instances of inherited systems that implement custom features like rendering of a 3D structure represented by connected root segments.

Note that the concept of the system does not mandate a mapping to structural components. Some systems may be designed to contain variables, such as length and 3D coordinates, for keeping track of the structural growth of a specific part of the plant, but the systems are no different from other systems without such variables and there is no technical distinction between structural and non-structural components in terms of data types. Any implication of the system is entirely determined by the behavior of variables declared within the system.

#### 2.2.1 Context

Every system has an internal variable named `context` which is a non-state variable referencing to an instance of `Context` system. Context serves two critical purposes in simulation: time and configuration management.

~~~
@system Context **begin**
  context ∼ ::Nothing
  config ∼ ::Config(override)
  clock(config) ∼ ::Clock
**end**
~~~

`Clock` system referred by `clock` variable in `Context` keeps track of a few time-related variables, namely `init`, `step`, `time`, and `tick`. `time` is a variable declared as `advance` which is essentially an `accumulate` variable tailored for keeping time of simulation. `time` accumulates from an initial value `init` by an interval value `step` for each update call that governs an overall time frame of the simulation. Without custom configuration, `time` starts from 0 hour and increases by 1-hour interval that assumes hourly simulation of crop models. `tick` is another time variable keeping the same time frame by just counting the number of updates done so far. Thus `context.clock.time` or `context.clock.tick` is often used as an index of data frame storing simulation output and as x-axis of a plot visualizing such output. Most of the time, a single instance of context is shared between multiple systems under a homogeneous time frame. Adopting heterogeneous time frames for a certain group of systems is then allowed by assigning an instance of a context with independent clock configuration to them. `Config` referred by `config` variable in `Context` is not a system, but a configuration object structured as a nested dictionary or hash table to store user-defined parameter value as a triplet of *system – variable – value*. A parameter value specified in configuration is plugged into a `preserve` variable with `parameter` tag during initialization of the variable. The framework supports a couple of handy operations around configuration objects to facilitate a large-scale simulation, *i*.*e*., in temporal and spatial domains. A new configuration object may be created by merging two or more existing configurations. Since a parameter value referring to the same variable of a certain system would be overridden by a value from a later configuration, Cropbox encourages organizing a set of small pieces of configuration to assemble a large one. Helper functions are also provided to populate multiple configuration objects from a range of values assigned for a specific parameter. For convenience, @config macro provides operator-based syntax to support complex operations like merging and expansion of configuration in a relatively simple statement.

#### 2.2.2 Controller

An instance of context and configuration provided to a new instance of system usually comes from a parent system that owns a variable referring to the system and thus initiates an instantiation of it. However, since there would be no parent system for the first system getting instantiated, context and configuration need to be supplied elsewhere.

~~~
@system System **begin**
  context ∼ ::Context(override)
  config(c = context) => c.config ∼ ::Config
**end**
~~~

`Controller` is a built-in mixin to handle such initiation issues by creating an instance of `Context` by itself. `Config` object is overridden by a keyword argument (`config`) of system constructors such as instance() and functions internally make use of it such as simulate() and visualize(). Therefore at least one, and usually only one, system is designated to possess `Controller` as the last element of its mixin list. The system augmented with `Controller` is what model users expect to primarily interact with in their workflow.

~~~
@system Controller **begin**
  config ∼ ::Config(override)
  context(config) ∼ ::Context(context)
**end**
~~~

### 2.3 Implementation

The Cropbox modeling framework is implemented as a package named Cropbox.jl^1^ written in Julia programming language [Bezanson et al., 2017]. Julia is a recently emerging language with a strong focus on scientific computing where often demands high performance while also benefits from features of dynamic programming languages such as Python. Especially, dynamic code expansion using macros allowed implementation of the framework in a domain-specific language with semantics covering the concept of system and variable in a surface syntax readily supported by Julia language.

A Cropbox *system* is defined with @system macro followed by a block containing variable declarations with the syntax described above. When this macro is expanded, the framework generates code for defining a new `struct` for the system and several internal functions around it. Each variable becomes a field in the `struct` wrapped with corresponding `State` type equipped with a variable-specific code generation logic. The new `struct` is defined to be a subtype of `System` or an optional base type as specified in the declaration for inheritance purposes.

During macro expansion, the framework keeps track of all variables declared in the system and creates a dependency graph based on dependent variables listed in the declarations. A topological sorting algorithm is then applied to the dependency graph to delineate the order between variables. For instance, if variable `b` depends on variable `a`, update code for `a` must execute before updating `b` to prevent premature access to an old state of `a` from the previous time step. A determined variable ordering is used in code generation to properly lay down variable handling logic in system initialization and update functions.

Some variable types may rely on a more fine-grained control of update sequences. For example, `accumulate` variable has to calculate a new rate for the next time step by the end of the current time step while this new rate should not be reflected in the accumulated value in the current time step yet. These two operations, accumulate by a rate from the last time step and calculating a new rate for the next time step, are internally split into *main* and *post* steps which become two separate nodes in the dependency graph. Any reference to this variable makes two dependent edges to both nodes while *post* node has a dependent edge on *main* node. This deliberate separation between two nodes ensures that the calculation of the new rate takes place after finishing updates of all other variables relying on this variable. `bisect` variable uses a similar mechanism with two steps, *pre* and *main*, to form a loop implementing iterative updates of solution. For less common circumstances where per-system level control flow using steps is not enough, the framework provides per-context level hooks called stages to allow inserting code before or after updating all systems under the same context. `produce` variable relies on three stages, *pre, main*, and *post*, to instantiate a new system before any other variables can see it at the beginning of a time step while passing necessary information for instantiation to the next time step. Note that these technical details are not necessarily exposed to the model developers and are abstracted away thanks to the succinct form of declarative syntax.

Some parts of the Cropbox syntax are based on the integration with other packages and the standard Julia runtime environment. For example, measurement units are defined using @u_str macro (*e*.*g*. `u”m/s”`) provided by Unitful.jl^2^ package. Strings (docstrings) placed in front of the declaration of a variable or system are automatically transformed into documentation which becomes available through the standard Julia help prompt or Cropbox built-in `look()` function or @look macro for introspection.

### 2.4 Workflow

Cropbox provides several core functions to support a typical workflow for crop model simulations. simulate() function accepts a system generated with @system macro to run a dynamic simulation over multiple time steps until a certain stop condition is met while inspecting the current state of running systems for each time step to extract variables of interest into a tabular data frame.

~~~
simulate(Garlic.Model;
  config = Garlic.Examples.AoB.KM_2014_P2_SR0,
  stop = “calendar.count”,
)
~~~

Stop condition (`stop`) can be simply a total number of updates or time period suffixed with proper time units such as hour and day. If more sophisticated control is needed, a dedicated `flag` variable can be declared in the model and passed as a stop condition. For maximum flexibility, any function accepting an instance of a system to implement custom logic for checking stop conditions can be also used.

For models relying on external parameters, the configuration option (`config`) can accept either a single configuration object or a list of multiple configurations. In the latter case, multiple iterations of simulation for each configuration object will be conducted and the output will contain a result aggregated from each simulation. Multithreading can be enabled for maximizing performance on multi-core processors.

With an output stored in a data frame, which is a common format used by data science packages in Julia, users can proceed with further analysis and visualization on their own. Specifically, Cropbox provides visualize() function to allow plotting several different kinds of graphs based on the list of variables declared in the system. Name of the variable in a symbol (*e*.*g*. `:A_net`) or a full path of variable in a string with enclosing system names (*e*.*g*. `“LeafGasExchange.ModelC4MD.A_net”`) are passed as arguments to determine horizontal and vertical axes of the graph.

~~~
visualize(ModelC4MD, :Ci, [:A_net, :Ac, :Aj];
  config = ge_base,
  xstep = Weather => :CO2 => 10:10:500,
  kind = :line,
)
~~~

Multiple variables of similar units and scales can be specified via grouping option (`group`) to draw multiple lines of variables (*e*.*g*. `A_net` and `A_gross` at the same time) with distinct colors and labels for visual comparison.

In an interactive programming environment (*e*.*g*., Jupyter Notebook [Kluyver et al., 2016]), interactive exploration of plots can be facilitated with manipulate() function similar to visualize() but extended with a list of value ranges for parameters controlling the shape of graphs. An interactive user interface is rendered in the notebook for each parameter with a slider widget that users can freely change a parameter value. A change in the slider would trigger a new simulation running under the hood and a new plot redrawn on the fly.

~~~
manipulate(ModelC3BB, :T, :A_net;
  config = ge_base,
  parameters = (
    Weather => :PFD => 0:100:2000,
),
  xstep = Weather => :T_air => 0:1:45,
  group = Weather => :CO2 => [1000, 400, 250],
  xlim = (0, 45),
  ylim = (0, 35),
  kind = :line,
)
~~~

When an optimal set of parameters is needed, calibrate() function supports parameter fitting using a global optimization method provided by Black-BoxOptim.jl^3^ package. The optimization method attempts to find the best fit set of parameters under a given observation dataset and given boundaries of parameter values. The observation dataset should be in a similar structure to the output data frame where a mapping between variables for an existing observation and an estimation from the model exists. Variables can be categorized into *index* or *target* depending on their role in the mapping. Index variables become index columns when two datasets are joined for comparison and usually consist of time components in the case of dynamic simulation models. Target variables are non-index variables that are subject to the evaluation of fitness.

~~~
calibrate(ModelC3BB, obs, obs_configs;
  index = [
    :PARi => :PFD,
    :CO2S => :CO2,
    :RH_S => :RH,
    :Tair => :T_air,
    :Press => :P_air,
],
target = [:Photo => :A_net, :gs],
parameters = (
  StomataBallBerry => (;
     g0 = (0, 1),
    g1 = (0, 10),
),
),
)
~~~

For convenience, evaluate() function provides several evaluation metrics like RMSE (root mean square error) and Nash-Sutcliffe model efficiency (*a*.*k*.*a*., model efficiency (EF); Nash and Sutcliffe [1970]) using the same interface as `calibrate()`.

~~~
evaluate(obs, est;
  index = :measuring_date => :date,
  target = :bulb_dry_weight => :bulb_mass,
  metric = :ef,
)
~~~

All workflow functions are supported on the Jupyter notebook environment which is well suited for running existing models for explanatory research. For developing a model as a package organized with more than a single file, developers may prefer a conventional development environment based on a code editor and REPL (read-eval-print loop). Cropbox was developed to support an entire modeling and simulation workflow running under a terminal-oriented environment. For example, dive() function allows inspection of the current state of a system instance by navigating enclosed variables using keyboard shortcuts in REPL. Plots can be still rendered in a legible format using a set of text letters without heading to a notebook.

## 3 Applications

### 3.1 Coupled Leaf Gas-Exchange Model

A coupled gas-exchange model is an important building block of a crop model for a mechanistic assessment of CO_2_, water vapor, and energy exchanges under diverse environmental conditions. Our implementation of the coupled gas-exchange was based on an existing model written originally in Object Pascal/Delphi language for rose leaves [Kim and Lieth, 2003] and later translated to C++ to become part of a maize model (MAIZSIM; Kim et al. [2012]) and a garlic model [Hsiao et al., 2019]. The model was then reimplemented in Cropbox and recently used to compare the behavior of stomatal conductance sub-models integrated with a coupled gas-exchange model for C4 leaves [Yun et al., 2020] (Figure 2).

**Figure 2:**
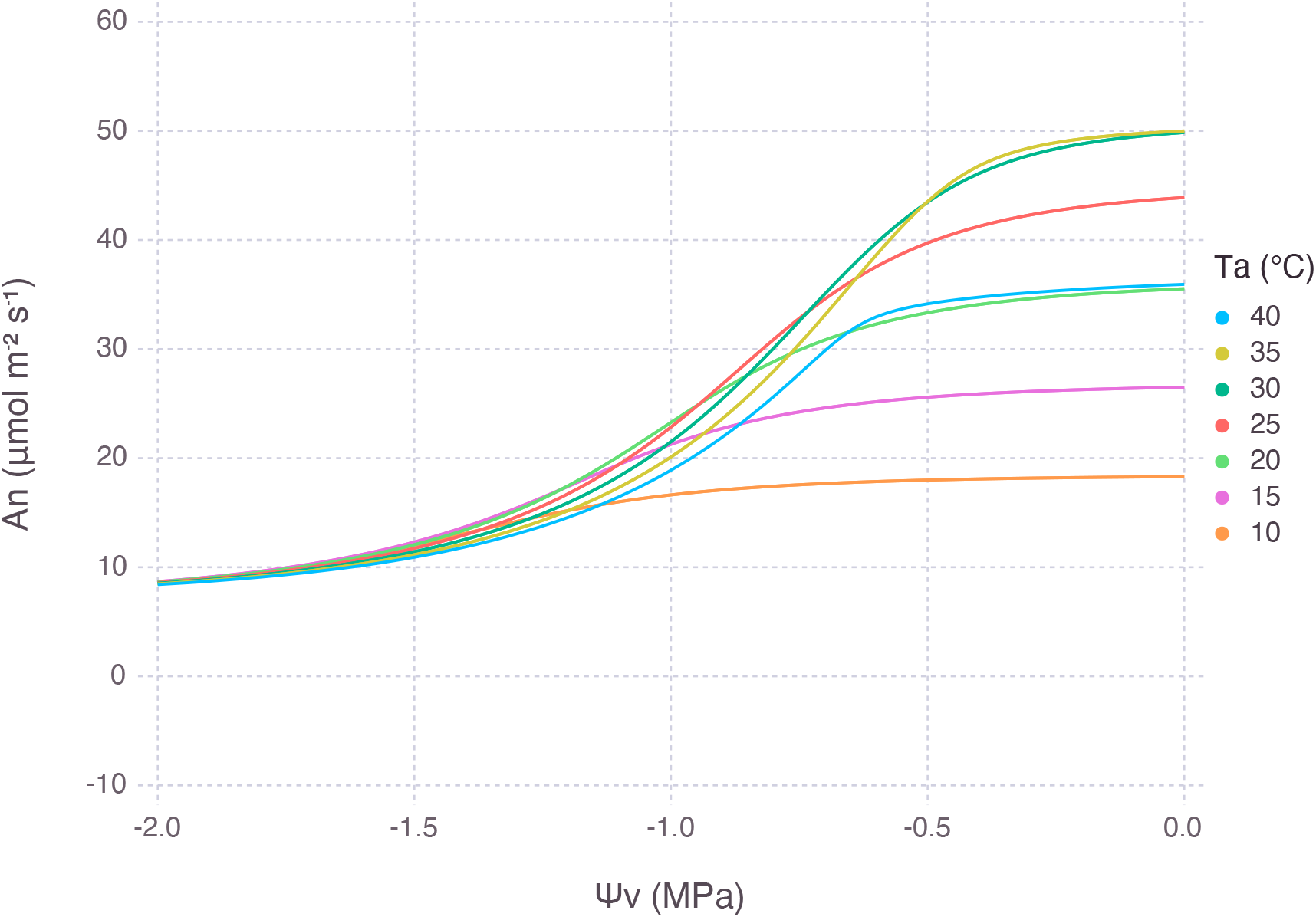
Net photosynthesis rate of a C_4_ leaf simulated by LeafGasExchange.jl under diverse combinations of air temperature (*T*_*a*_) and leaf water potential (Ψ_*v*_). The relative strength of water stress was dependent on temperature which was a primary factor determining a potential rate of photosynthesis.

The original model [Kim and Lieth, 2003] was coded in a monolithic class that contains all parameters and variables in a single location. Variables were updated via explicit calls to several functions representing sub-models such as stomatal conductance and energy balance. These functions were written carefully to calculate new values of variables in the correct execution order. There were iterative solvers implemented in two locations, one for obtaining intercellular CO_2_ (*C*_*i*_) and the other for adjusting leaf temperature. Tracking iterations and updating respective variables had to be coded explicitly inside the loop. Some parameters were stored in a separate structure to allow customization before running the model. Output variables were available via instance methods and a few of them were saved in a file after the simulation was done, but further analysis and visualization would require external tools.

The updated model reimplemented in Cropbox, released as LeafGasExchange.jl^4^ package, has a modular structure with more than 30 small systems representing a consolidated aspect of the gas-exchange process and its hierarchy. For example, three different stages of limitation in C_3_ biophysical photosynthesis were divided into separate systems, namely `C3c`, `C3j`, `C3p`, for representing enzyme-limited, electron transport-limited, triose phosphate-limited pathways, respectively [Sharkey, 2016]. Then `C3` system was composed with these components to provide net photosynthesis rate (`A_net`) and the likes by comparing rates calculated from different pathways. Variables were declared with explicit units such as μmol m^−2^ s^−1^ for net photosynthesis (`A_net`) and mol m^−2^ s^−1^ bar^−1^ for stomatal conductance (`g_s`). The difference in scaling between units like μmol and mol were automatically handled via implicit conversion triggered on variable updates.

A complex control structure introduced to implement an iterative solver in the original model was replaced by `bisect` variable. The bisection method is slow to converge but is guaranteed to find a solution if one exists within a given boundary, unlike faster methods occasionally failing under a certain condition such as low atmospheric CO_2_ concentration. In the case of solving the energy balance equation, the adoption of the bisection method allowed the removal of the approximation of leaf temperature relying on Taylor expansion which led to an intuitive code that looked exactly the same as the original equation (*R*_*n*_ = *H* + *λE*).

~~~
ΔT(R_net, H, *λ*E): temperature_adjustment => **begin**
  R_net == H + *λ*E
**end** ∼ bisect(lower = -5, upper = 5,
     u”K”, evalunit = u”W/m^2”)
~~~

Biochemical photosynthesis and stomatal conductance are coupled via relative humidity at the leaf surface (*H*_*s*_) which is solved based on an empirical relationship [Collatz et al., 1991]. Instead of manually describing a lengthy solution of the quadratic equation, we can use a `solve` variable with the equation to be solved specified inside the body. Then an identical symbolic solution will be automatically generated during macro expansion and its code will be used as if we have a `track` variable for the solution.

~~~
hs(g0, g1, gb, An, Cs, RH): humidity_surface => **begin**
  gs = g0 + g1*(An*hs/Cs)
  (hs - RH)*gb == (1 - hs)*gs
**end** ∼ solve(lower = 0, upper = 1)
~~~

Cropbox supports the selection of arbitrary output variables instead of a fixed set as in the original model that allowed model developers and users freely look into any variables of interest by running simulations with corresponding `target` variables. In the original model, adding new output variables requires modification of source code followed by recompilation which can easily interrupt exploratory workflow. With extended features and added usability, LeafGasExchange.jl (version 0.1.0) was written in 407 lines of Julia code which was 40% smaller compared to the comparable standalone model written in 680 lines of C++ code.

### 3.2 Process-based Crop Model for Garlic

Crop model is an integrated collection of multiple building blocks such as ontogenic development and structural growth with carbon assimilation and allocation. A process-based garlic crop model was developed based on the coupled gas-exchange model reviewed above with canopy-level scaling in conjunction with key phenological and morphological traits of the crop to describe a whole-plant growth [Hsiao et al., 2019]. The original model written in C++ was reimplemented in Julia using the Cropbox framework and released as Garlic.jl^5^ package [Yun et al., 2022] (Figure 3).

**Figure 3:**
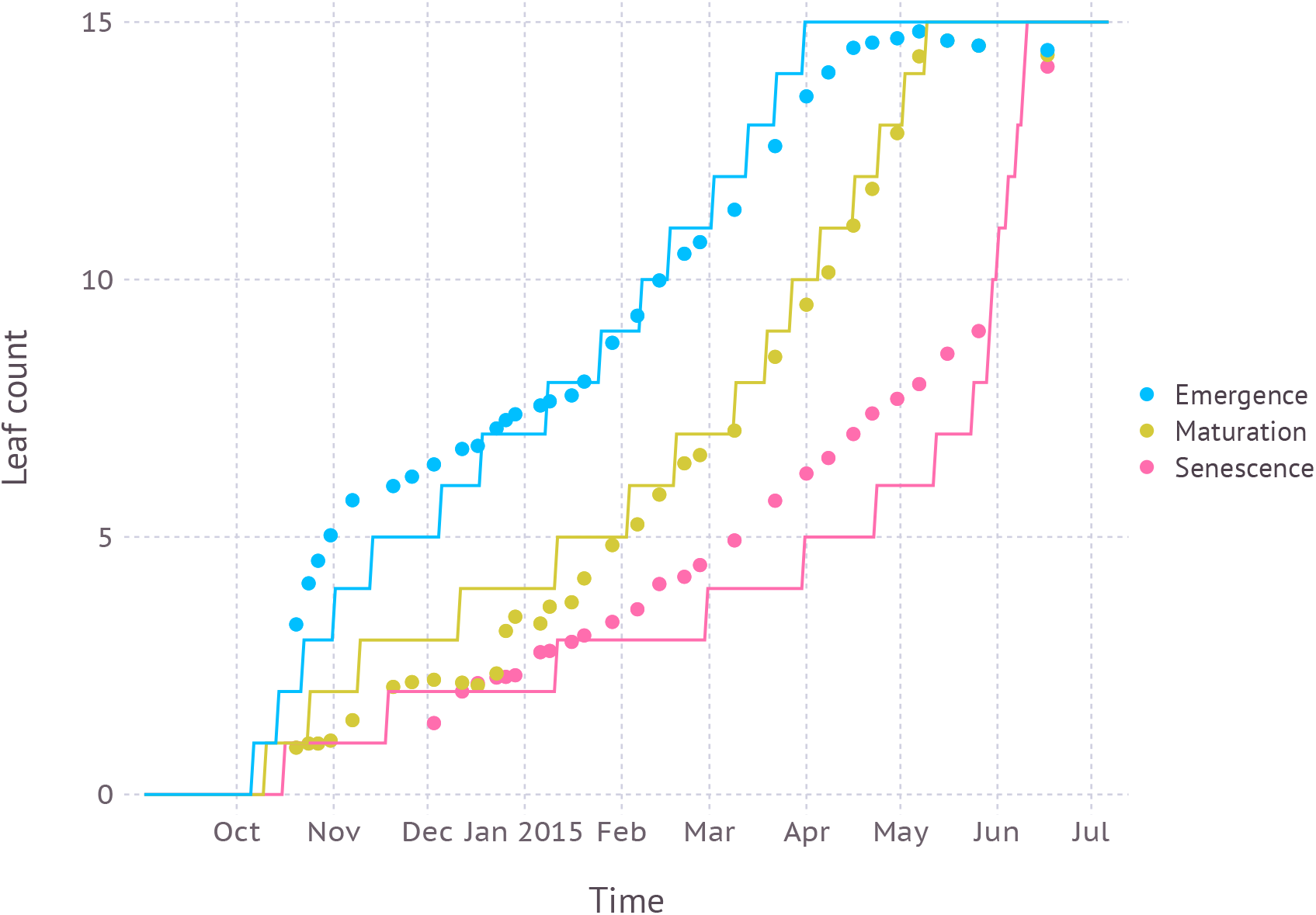
Leaf development simulation for garlic (*Allium sativum* ‘Shantang Purple’) ran with Garlic.jl and compared with observed field data. This figure reproduces Fig. 3A from Hsiao et al. [2019] using the same dataset and parameters.

The original model was composed of 16 classes representing plant structure and relevant biophysical processes. In the reimplemented model, they were expanded into more fine-grained 63 systems to cover the same functionalities. For example, a single class implementing coupled gas-exchange model was translated into 14 systems as illustrated in the previous section. Systems were placed under directories named **morphology, phenology, physiology, atmosphere**, and **rhizosphere** according to their roles in the model.

For **phenology**, a single class for tracking development in the original model was split into a set of small systems by consolidating each phenology stage into a separate system. Such reorganization of code into a smaller unit increases the cohesion of each system by grouping interdependent variables while enforcing a boundary between unrelated variables. A fine-grained organization of systems also entails a similar hierarchy of configuration which effectively groups enclosed parameters into more specific categories with better identifiability.

**morphology** contains systems describing the physical structure of a plant where the leaf was given most details due to its importance in light capture and gas-exchange dictating the overall growth of a plant. Each individual leaf is described by a single instance of `Leaf` system with its own tracking of elongation and life-cycle according to thermal time accumulation. An instance of enclosing nodal unit (*i*.*e*., phytomer) is dynamically created and added runtime via `produce` variable. Aside from leaves, systems for other structures and processes are static in the sense that no addition or removal of instances occurs once systems are initialized at the beginning of the simulation, like a static architecture of typical crop models.

Some variables need to aggregate values from multiple instances of a dynamic system. For example, total leaf area in `Area` mixin system in **physiology** would be a sum of leaf area from individual leaf instances inside nodal units. Such aggregate variables can be declared with an aggregate syntax such as summation using the index key on `produce` variable.

~~~
leaf_area(x = NU[“*”].leaf.area) => **begin**
  sum(x)
**end** ∼ track(u”cm^2”)
~~~

`leaf_area` variable is declared in `Area` mixin for calculating total leaf area by summing up individual leaf areas from all nodal units (`NU[“*”]`). Each nodal unit has `leaf` which is an instance of `Leaf` system and each leaf has a variable `area` for tracking leaf area growth over time. sum() is a standard Julia function for calculating the sum of all elements in a collection. `NU[“*”].leaf.area` returns a collection of leaf area from individual leaves.

**physiology** contains several mixins for `Plant` system which encompasses all other systems in the package. Notable mixins include `Area` for tracking total leaf area, `Mass` for tracking biomass of each plant structure, `Count` for leaf counting. `Photosynthesis` is a mixin for scaling up leaf-level `GasExchange` to canopy-level by incorporating sunlit and shaded leaf approach with canopy radiation calculated via `Radiation` mixin. The `Garlic.Model` system which is a primary system that users interact with is indeed `Plant` with `Controller` added for making the system instantiable.

With the original C++ model, dedicated Python scripts had to be written for running multiple simulations with template-based configuration files and for aggregating results from parsing output files. With Cropbox, this workflow was streamlined as the use of simulate() and @config provides a uniform interface regardless of model implementations so that the new garlic model no longer had to rely on custom script files. Albeit more features, Garlic.jl (version 0.1.16) was written in 1987 lines of Julia code which was smaller than 2750 lines of C++ code written for the original garlic model (version 0.1.10).

### 3.3 3D Root Architecture Model

While the Cropbox modeling framework was primarily designed for developing process-based crop simulation models with a static structure as seen by the examples above, the framework itself does not preclude models with a more dynamic structure which is often demonstrated by functional-structural plant modeling (FSPM) approaches [Vos et al., 2010, Soualiou et al., 2021]. A major difference with such structure-oriented models compared to the conventional crop simulation models is that the former often needs to deal with a sheer amount of structural components that have to be dynamically produced and interconnected for describing a specific aspect of plant growth, which is hardly captured by empirical models relying on a simpler structure approximation. As structures are dynamically generated during simulation, the framework must provide means to handle the addition of a new instance of a system at a proper timing compatible with how other variables work as briefly explained with the internals of `produce` variable relying on three stages. Support for aggregation via a special syntax also helped manage a large number of systems in a consistent way.

Implementation of the 3D root structure growth model was based on an algorithm proposed by CRootBox [Schnepf et al., 2018]. With the existence of multiple root types depending on the branching order, a strain of each root type consists of multiple root segments. In our implementation of the algorithm as a part of CropRootBox.jl^6^, the root type corresponds to a system and the root segment is then represented by an instance of a certain system. Like the dynamic production of individual leaves in the garlic model, each root segment is produced from another root segment when the accumulated length of connected segments exceeds a certain threshold specified by a parameter. For randomness in the growth, parameter values are given with standard deviations (*i*.*e*. `:S => :a => 1.0` *±* `0.1`) and the actual values are sampled from corresponding normal distributions on initialization. As new root segments can be added on two occasions, axial growth and lateral branching, two separate `produce` variables were used. For interoperability between different root types, ‘BaseRoot‘ which contains all necessary parameters and state variables as well as a branching transition table is inherited by concrete subtypes of each root such as `PrimaryRoot` and `FirstOrderLateralRoot` systems.

The current position of a root segment is calculated with a chain of transformation matrices. The local transformation matrix is determined by the length of the elongated segment (`l`) and branching angle from the parent segment, namely axial angle *α* and radial angle *β*. For a current position vector (`cp`) in global coordinate system, global transformation matrix (`RT1`) is obtained by multiplication of local transformation matrix (`RT`) and previous global transformation matrix of parent segment (`RT0`). A reference to the variable holding global transformation matrix (`RT1`) is passed down to subsequent systems by overriding a `track` variable itself pointing to the parent transformation matrix (`RT0`) rather than by mere copying of matrix values in order to ensure any changes made in parent system, such as translation vector change due to root elongation, automatically propagates down to child systems so that global transformation matrices (`RT1`) in child systems receive proper updates for each time step.

~~~
RT0: parent_transformation ∼ track::Transformation(
   override)
RT(nounit(l), *α, β*): local_transformation => **begin**
  T = Translation(0, 0, -l)
  R = RotZX(*β, α*) |> LinearMap
  R ° T
**end** ∼ track::Transformation
RT1(RT0, RT): global_transformation => **begin**
  RT0 ° RT
**end** ∼ track::Transformation
cp(RT1): current_position => **begin**
  RT1(Point3f(0, 0, 0))
**end** ∼ track::Point3f
~~~

Once the simulation is complete, a network of more than thousands of root segments could be produced as a result. For rendering root structure in 3D space, a custom Julia function was written to collect a list of meshes generated from each root segment by recursively visiting systems with `Rendering` mixin which worked as a trait without further variable declarations. A collected list of 3D mesh can be used to make an interactive 3D visualization directly launched from a Julia session or to export to a VTK file for further processing with external programs (Figure 4).

**Figure 4:**
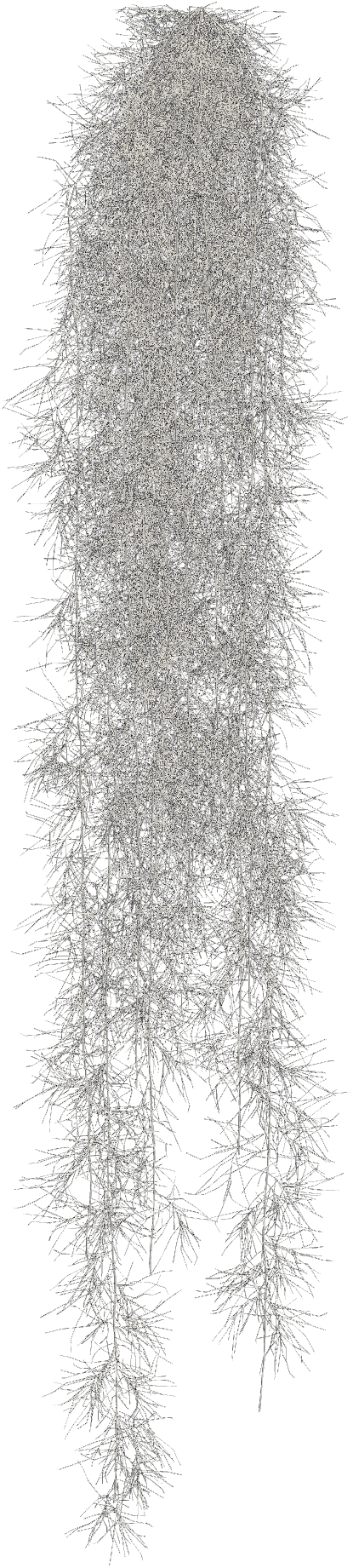
Rendering of 3D root structure simulated by CropRootBox.jl for switchgrass (*Panicum virgatum*). Three root types were modeled by corresponding systems and a large number of instances of these small root segments were dynamically generated during simulation via `produce` variable.

CropRootBox.jl (version 0.1.2) was written in 356 lines of Julia code. For a naive comparison, CRootBox^7^ (as of Nov 6, 2019) was written in 4610 lines of C++ code with additional Python bindings but came with more complete built-in features like segment analysis. Note that CropRootBox.jl was never intended to be a full reimplementation of CRootBox, but built as a proof of concept for applying the FSPM approach within Cropbox framework. For this purpose, we believe CropRootBox.jl showed a potential to fill the gap between conventional crop modeling and FSPM within the same framework. Future work will be focused on supporting custom functions with less boilerplate code by providing a more intuitive interface for matching and aggregating multiple instances of systems stored in `provide` variable.

## 4 Discussion

### 4.1 Variable-centric Modeling

Declarative syntax of Cropbox forces a model to be represented in terms of variables. Each variable is explicitly declared with an intended behavior (state) and associated metadata, such as a unit of measurement and range of value. Since the variable is subject to performing a small unit of task with a predefined set of operations, any chance of logic errors and human mistakes permeating through model code is greatly reduced. Such a declarative programming style is effective to prevent side effects like an unintended mutation of variables and incautious change of control flows [Muetzelfeldt, 2004, Rahman et al., 2004]. The correct order of execution and thus the precise scope of side effects should be automatically resolved during the code generation stage when the model specification is lowered to regular Julia code. With a carefully chosen set of options for each type of variable, model developers refrain from taking direct control over how the final code is laid out, producing more robust code.

Once incorporated into a dependency graph that describes the entire model, the variables can be easily accessed via a standard interface provided by the framework. For example, any declared variables can be referred to by their names to be a target of built-in functions like `simulate()` and `visualize()` when traditional models are often programmed in a way requiring code changes to expose internal state variables. Since the meaning of the variable and its relationship to others are explicitly declared, visualizing a dependency graph of the model becomes straightforward and provides a useful insight for understanding the complex structure of the model (Figure 1(b)).

In practice, explicit declaration of individual variables encourages a clear separation between variables interrelated with a certain mathematical meaning. Some intermediate variables, otherwise would have disappeared in the middle of the equation, would now show up as self-contained variables with distinct names and exact units. Making smaller computation units has been suggested as a good software engineering practice for improving code documentation and testing [Holzworth et al., 2015].

### 4.2 Unit Handling

Unit of measurement (*e*.*g*. `u”`*μ*`mol/m^2/s”`) attached to the variable is not merely cosmetic, but actively participates in automatic unit conversion and validation by a seamless integration of Unitful.jl package. The framework can recognize any misuse of units in the model when the calculated unit of a variable does not match the specified unit. In our experience, this feature alone greatly reduced logical errors and mistakes during model development even before having to inspect the output of the model simulation. It also eliminates the need for unit scaling factors in most cases when the variables are defined in compatible units. For example, a variable of `u”cm”` unit and another variable of `u”m”` can be used interchangeably as dependent variables in the declaration of others without a manual scaling factor like 100. The calculated unit will be then always converted to the unit declared with the variable in the end. Such behavior helps simplify equations shared across different sub-models and also catches incorrect scaling of empirical equations due to implicit assumptions on the units.

### 4.3 Runtime Performance

Julia is a dynamic language but takes advantage of just-in-time (JIT) compilation to provide runtime performance close to statically-typed languages [Bezanson et al., 2017]. Julia code running for the first time in the current session typically needs to be compiled which takes some time and causes a delay in interactive use. Once the code is compiled, any reuse of the same code, *i*.*e*. calling the same function again, runs significantly faster. For qualitative comparison, compiled Julia code often runs much faster than native Python code and can come close to the performance of C and Fortran when properly optimized [Sells, 2020].

We originally developed a prototype of Cropbox in Python to take advantage of metaclass and annotation syntax. However, after initial testing, we switched to Julia because of its higher performance with more flexible metaprogramming features. In addition, Cropbox also supports multi-core CPU for automatically running simulations in parallel for scalable performance when multiple configurations, *e*.*g*. representing multiple years and sites, are configured.

### 4.4 Interactive Workflow

With that, Cropbox framework is currently more aligned with the goal of providing adequate runtime performance for exploratory interactive use rather than achieving the highest possible speed for finishing simulations. Model developers often work in an interactive development environment like Julia REPL (Read-Eval-Print Loop) and Jupyter Notebook where incremental model development and simulation is possible.

Let’s say a developer is building a model for biomass growth. An initial model would be bare-bones like having an `accumulate` variable with a constant rate fixed in `preserve`. After writing the said model in two-lines of `System`, the developer can quickly test the model by running `simulate()` and/or `visualize()` on it. In the next line in the prompt, a refined version of the model, now with a proper rate variable defined in `track`, would be typed in. Evaluate the model. Perhaps the next revision would include a few parameters in `preserve` to replace some constant numbers in the rate variable.

Cropbox supports such an interactive workflow of model development in one continuously running session, without needing to switch back and forth between multiple separate sessions for building the model (*e*.*g*. in C), running the model (*e*.*g*. in a shell script), evaluating the model output (*e*.*g*. in Python), modifying the model (*e*.*g*. back in C) and repeating the tasks. This level of interactivity is usually not possible with statically compiled crop models.

The support of interactive workflow was also found to be a critical component for developing effective educational resources (Figure 5). We have extensively used Cropbox for teaching a graduate-level plant modeling course and an upper-level undergraduate plant ecophysiology course, as well as organizing workshops on crop modeling targeted towards horticultural researchers [Yun et al., 2021]. Most of the audience had no prior experience with Julia but successfully finished the course by focusing on the specific use of Cropbox framework.

**Figure 5:**
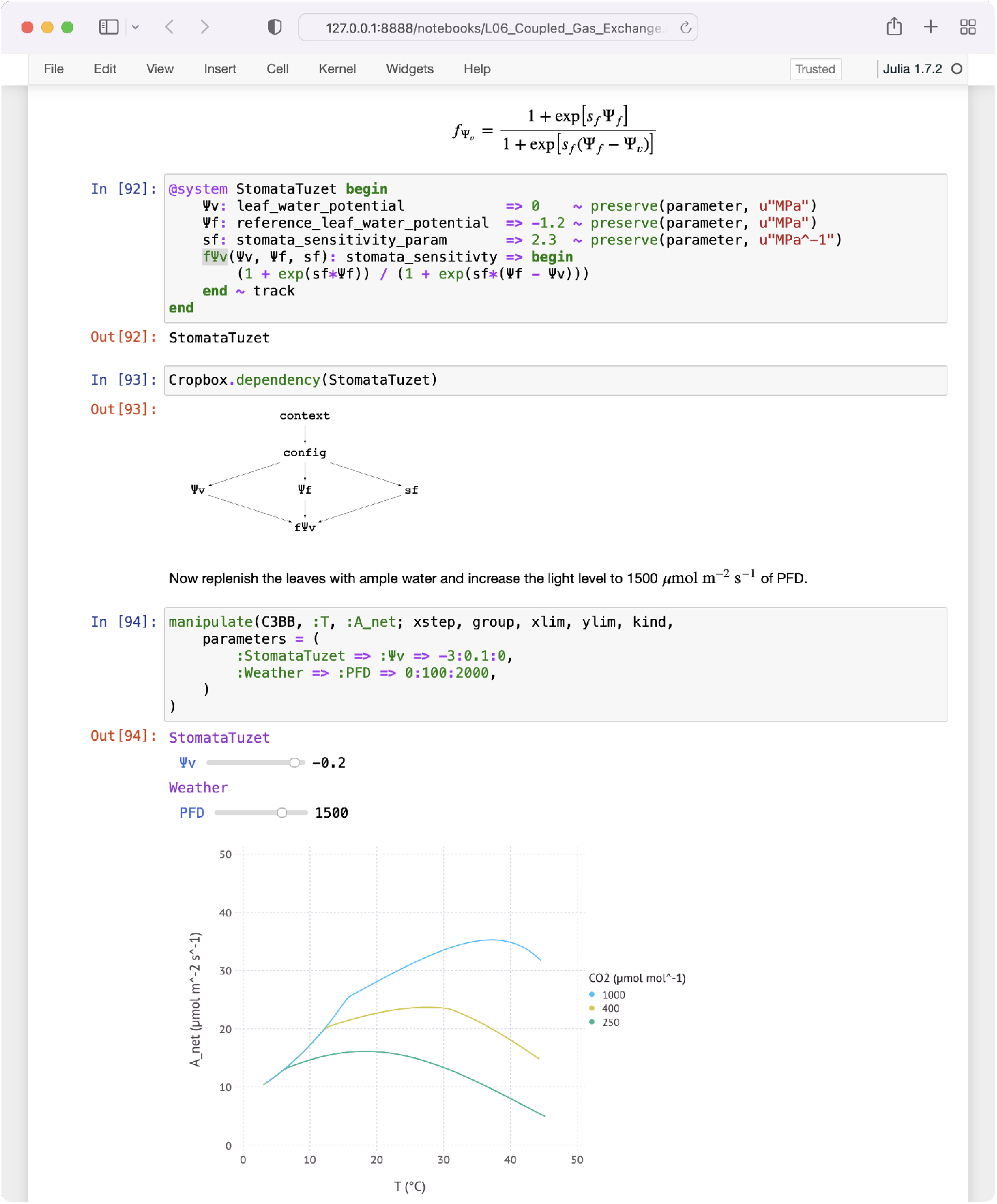
An example of Jupyter Notebook explaining how leaf-level gas exchange could respond to a varying degree of water stress and light level. `manipulate()` function creates a user interface for interactive plot. Whenever the user adjusts a slider widget to change one of the selected model parameters, the plot will be dynamically updated with a new simulation result.

### 4.5 Framework Extension

In addition to the currently supported 19 types of state variables, there is an opportunity open for adding new types to the framework as needed. Many of the current state types were derived during the rewrite of existing models in other languages. For example, `bisect` was added to implement the bisection particularly needed for finding a root in the coupled gas-exchange model. The original model had the root-finding algorithm directly embedded in the model code with no separation of computational logic for other variables, leading to a complicated mixture of code. Cropbox reduced the code complexity by providing a new layer of abstraction for the root-finding hidden by the `bisect` state specifically designed for the purpose. Any common design pattern identified for a particular logic reusable by other models would be a good candidate for implementing a new type of state.

Any `System` declared in Cropbox is translated into a regular Julia `struct` with associated internal functions automatically generated for the integration with built-in workflow functions like simulate(). For advanced applications, model developers can design a custom interface for extending features beyond the framework currently provides. For example, CropRootBox.jl implemented its own render() function for visualizing 3D geometries of the root architecture (Figure 4). The render() function iterates over the network of `RootSegment` systems and constructs a collection of cuboid mesh corresponding to each segment which is finally rendered by the external visualization package Makie.jl [Danisch and Krumbiegel, 2021].

Other features extending the framework currently in development include, but not limited to, surrogate() function to make a surrogate machine-learning model trained by the results of simulation from existing `System`, rasterize() function to extract a grid of `Config` objects from geospatial raster datasets, calibrate() function to use a custom optimization method wrapped in a user-supplied callback, and panelize() function to create an interactive dash-board extending the existing manipulate() for a full-fledged user interface for controlling model simulation and analysis.

To support Cropbox-specific workflow in a more traditional development environment, editor plugins could be also developed for syntax highlighting, auto-completion, and code visualization using dependency graphs. We could even think about alternative backends for the framework, for example, emitting host code in languages other than Julia for the case when interoperability with the existing system is highly prioritized.

## 5 Conclusions

We introduce Cropbox, a novel modeling framework that supports various aspects of crop modeling in a unique yet concise style. Building a crop model can be easily riddled with technical details looking trivial at first, but later becoming major obstacles that hamper the whole development process, especially when implementing models from scratch without relying on an established framework. Cropbox adopts a declarative approach using a domain-specific language (DSL) to reduce the technical burdens and let the model developers focus on a high-level abstraction with the meaning of the variable and its relation to others, rather than tinkering with low-level implementation details. The syntax of the surface language is deliberately constrained to avoid unintended side effects caused by common mistakes, but the architecture is still open to extension. We anticipate that Cropbox will become a useful tool for crop modelers with their development and application of crop models in ways that have yet to be seen. We also hope Cropbox will become an agent that blurs the lines between model users and model builders, with fewer technical hurdles.

## 8 Contributions by the authors

**Kyungdahm Yun**: Conceptualization, Methodology, Software, Data curation, Investigation, Visualization, Writing - Original draft preparation. **Soo-Hyung Kim**: Conceptualization, Resources, Writing - Review & Editing, Supervision, Project administration, Funding acquisition.

## Footnotes

1 https://github.com/cropbox/Cropbox.jl

2 https://github.com/PainterQubits/Unitful.jl

3 https://github.com/robertfeldt/BlackBoxOptim.jl

4 https://github.com/cropbox/LeafGasExchange.jl

5 https://github.com/cropbox/Garlic.jl

6 https://github.com/cropbox/CropRootBox.jl

7 https://github.com/Plant-Root-Soil-Interactions-Modelling/CRootBox

## References

B. Acock and V. R. Reddy. Designing an object-oriented structure for crop models. Ecological Modelling, 94(1):33–44, 1997.

M. Adam, M. Corbeels, P. A. Leffelaar, H. van Keulen, J. Wery, and F. Ewert. Building crop models within different crop modelling frameworks. Agricultural Systems, 113:57–63, 2012.

R. M. Argent. An overview of model integration for environmental application - components, frameworks and semantics. Environmental Modelling and Software, 19(3):219–234, 2004.

N. Athanasiadis and F. Villa. A roadmap to domain specific programming languages for environmental modeling. In ACM Workshop on Domain-Specific Modeling, pages 27–32, 2013.

B. N. Bailey. Helios: a scalable 3D plant and environmental biophysical modeling framework. Frontiers in Plant Science, 10:1185, 2019.

J. Benz, R. Hoch, and T. Legović. ECOBAS — modelling and documentation. Ecological Modelling, 38(1-3):3–15, 2001.

J. E. Bergez, P. Chabrier, C. Gary, M. H. Jeuffroy, D. Makowski, G. Quesnel, E. Ramat, H. Raynal, N. Rousse, D. Wallach, P. Debaeke, P. Durand, M. Duru, J. Dury, P. Faverdin, C. Gascuel-Odoux, and F. Garcia. An open platform to build, evaluate and simulate integrated models of farming and agro-ecosystems. Environmental Modelling and Software, 39(C):39–49, 2013.

J. E. Bergez, H. Raynal, M. Launay, N. Beaudoin, E. Casellas, J. Caubel, P. Chabrier, E. Coucheney, J. Dury, I. G. de Cortazar-Atauri, E. Justes, B. Mary, D. Ripoche, and F. Ruget. Evolution of the STICS crop model to tackle new environmental issues: New formalisms and integration in the modelling and simulation platform RECORD. Environmental Modelling and Software, 62(C):370–384, 2014.

J. Bezanson, A. Edelman, S. Karpinski, and V. B. Shah. Julia: a fresh approach to numerical computing. SIAM Review, 59(1):65–98, 2017.

G. Bracha and W. Cook. Mixin-based inheritance. SIGPLAN Notices, 25(10): 303–311, 1990.

R. D. Brennan and M. Y. Silberberg. The System/360 continuous system modeling program. Simulation, 11(6):301–308, 1968.

H. E. Brown, N. I. Huth, D. P. Holzworth, E. I. Teixeira, R. F. Zyskowski, J. N. G. Hargreaves, and D. J. Moot. Plant Modelling Framework: Software for building and running crop models on the APSIM platform. Environmental Modelling and Software, 62:385–398, 2014.

R. E. M. Caskie and R. E. A. Mason. Some Design Features Of Continuous System Modelling Program III. INFOR: Information Systems and Operational Research, 11(2):125–139, 1972.

G. Collatz, J. Ball, C. Grivet, and J. Berry. Physiological and environmental regulation of stomatal conductance, photosynthesis and transpiration: a model that includes a laminar boundary layer. Agricultural and Forest Meteorology, 54:107–136, 1991.

R. Costanza and S. Gottlieb. Modelling ecological and economic systems with STELLA: Part II. Ecological Modelling, 112(2-3):81–84, 1998.

R. Costanza and A. Voinov. Modeling ecological and economic systems with STELLA: Part III. Ecological Modelling, 143(1-2):1–7, 2001.

R. Costanza, D. Duplisea, and U. Kautsky. Ecological modelling and economic systems with STELLA. Ecological Modelling, 110(1):1–4, 1998.

S. Danisch and J. Krumbiegel. Makie.jl: Flexible high-performance data visualization for julia. Journal of Open Source Software, 6(65):3349, 2021.

O. David, J. C. Ascough II, W. Lloyd, T. R. Green, K. W. Rojas, G. H. Leavesley, and L. R. Ahuja. A software engineering perspective on environmental modeling framework design: The Object Modeling System. Environmental Modelling and Software, 39(c):201–213, 2013.

A. de Wit, H. Boogaard, D. Fumagalli, S. J. C. Janssen, R. Knapen, D. van Kraalingen, I. Supit, R. van der Wijngaart, and K. van Diepen. 25 years of the WOFOST cropping systems model. Agricultural Systems, 168:154–167, 2019.

M. Donatelli, G. Bellocchi, and L. Carlini. Sharing knowledge via software components: Models on reference evapotranspiration. European Journal of Agronomy, 24(2):186–192, 2006.

J. W. Forrester. Industrial Dynamics. M.I.T. Press, Cambridge, MA, 1961.

M. Henke, W. Kurth, and G. H. Buck-Sorlin. FSPM-P: towards a general functional-structural plant model for robust and comprehensive model development. Frontiers of Computer Science, 10(6):1103–1117, 2016.

C. Hillyer, J. Bolte, F. van Evert, and A. Lamaker. The ModCom modular simulation system. European Journal of Agronomy, 18(3-4):333–343, 2003.

N. Holst. A universal simulator for ecological models. Ecological Informatics, 13: 70–76, 2013.

D. P. Holzworth, N. I. Huth, and P. G. de Voil. Simplifying environmental model reuse. Environmental Modelling and Software, 25(2):269–275, 2010.

D. P. Holzworth, N. I. Huth, P. G. deVoil, E. J. Zurcher, N. I. Herrmann, G. McLean, K. Chenu, E. J. van Oosterom, V. Snow, C. Murphy, A. D. Moore, H. E. Brown, J. P. M. Whish, S. Verrall, J. Fainges, L. W. Bell, A. S. Peake, P. L. Poulton, Z. Hochman, P. J. Thorburn, D. S. Gaydon, N. P. Dalgliesh, D. Rodriguez, H. Cox, S. Chapman, A. Doherty, E. Teixeira, J. Sharp, R. Cichota, I. Vogeler, F. Y. Li, E. Wang, G. L. Hammer, M. J. Robertson, J. P. Dimes, A. M. Whitbread, J. Hunt, H. van Rees, T. McClelland, P. S. Carberry, J. N. G. Hargreaves, N. MacLeod, C. McDonald, J. Harsdorf, S. Wedgwood, and B. A. Keating. APSIM - Evolution towards a new generation of agricultural systems simulation. Environmental Modelling and Software, 62: 327–350, 2014.

D. P. Holzworth, V. Snow, S. J. C. Janssen, I. N. Athanasiadis, M. Donatelli, G. Hoogenboom, J. W. White, and P. Thorburn. Agricultural production systems modelling and software: Current status and future prospects. Environmental Modelling and Software, 72:276–286, 2015.

D. P. Holzworth, N. I. Huth, J. Fainges, H. E. Brown, E. Zurcher, R. Cichota, S. Verrall, N. I. Herrmann, B. Zheng, and V. Snow. APSIM Next Generation: Overcoming challenges in modernising a farming systems model. Environmental Modelling and Software, 103:43–51, 2018.

J. Hsiao, K. Yun, K. H. Moon, and S.-H. Kim. A process-based model for leaf development and growth in hardneck garlic (Allium sativum). Annals of Botany, 124(7):1143–1160, 2019.

S. J. C. Janssen, C. H. Porter, A. D. Moore, I. N. Athanasiadis, I. Foster, J. W. Jones, and J. M. Antle. Towards a new generation of agricultural system data, models and knowledge products: Information and communication technology. Agricultural Systems, 155(C):200–212, 2017.

J. W. Jones, B. A. Keating, and C. H. Porter. Approaches to modular model development. Agricultural Systems, 70(2-3):421–443, 2001.

J. W. Jones, G. Hoogenboom, C. H. Porter, K. J. Boote, W. D. Batchelor, W. D. Batchelor, L. A. Hunt, P. W. Wilkens, U. Singh, A. J. Gijsman, and J. T. Ritchie. The DSSAT cropping system model. European Journal of Agronomy, 18(3-4):235–265, 2003.

S.-H. Kim and J. H. Lieth. A coupled model of photosynthesis, stomatal conductance and transpiration for a rose leaf (Rosa hybrida l.). Annals of Botany, 91(7):771–781, 2003.

S.-H. Kim, Y. Yang, D. Timlin, D. Fleisher, A. Dathe, V. Reddy, and K. Staver. Modeling temperature responses of leaf growth, development, and biomass in maize with maizsim. Agronomy Journal, 104(6):1523–1537, 2012.

T. Kluyver, B. Ragan-Kelley, F. Pérez, B. Granger, M. Bussonnier, J. Frederic, K. Kelley, J. Hamrick, J. Grout, S. Corlay, P. Ivanov, D. Avila, S. Abdalla, C. Willing, and J. development team. Jupyter notebooks – a publishing format for reproducible computational workflows. In Positioning and Power in Academic Publishing: Players, Agents and Agendas, pages 87–90, 2016.

D. Kneis. A lightweight framework for rapid development of object-based hydrological model engines. Environmental Modelling and Software, 68(7): 110–121, 2015.

M. Lang. yggdrasil: a Python package for integrating computational models across languages and scales. in silico Plants, 1(1):diz001, 2019.

H. Lemmon and N. Chuk. Object-oriented design of a cotton crop model. Ecological Modelling, 94(1):45–51, 1997.

R. M Keller and J. L Dungan. Meta-modeling: a knowledge-based approach to facilitating process model construction and reuse. Ecological Modelling, 119 (2-3):89–116, 1999.

A. Marshall-Colon, S. P. Long, D. K. Allen, G. Allen, D. A. Beard, B. Benes, S. von Caemmerer, A. J. Christensen, D. J. Cox, J. C. Hart, P. M. Hirst, K. Kannan, D. S. Katz, J. P. Lynch, A. J. Millar, B. Panneerselvam, N. D. Price, P. Prusinkiewicz, D. Raila, R. G. Shekar, S. Shrivastava, D. Shukla, V. Srinivasan, M. Stitt, M. J. Turk, E. O. Voit, Y. Wang, X. Yin, and X. Zhu. Crops In Silico: Generating virtual crops using an integrative and multi-scale modeling platform. Frontiers in plant science, 8(786), 2017.

C. A. Midingoyi, C. Pradal, I. N. Athanasiadis, M. Donatelli, A. Enders, D. Fumagalli, F. Garcia, D. P. Holzworth, G. Hoogenboom, C. Porter, H. Raynal, P. Thorburn, and P. Martre. Reuse of process-based models: automatic transformation into many programming languages and simulation platforms. in silico Plants, 2(1):diaa007, 2020.

C. A. Midingoyi, C. Pradal, A. Enders, D. Fumagalli, H. Raynal, M. Donatelli, I. N. Athanasiadis, C. Porter, G. Hoogenboom, D. Holzworth, F. Garcia, P. Thorburn, and P. Martre. Crop2ML: An open-source multi-language modeling framework for the exchange and reuse of crop model components. Environmental Modelling and Software, 142:105055, 2021.

R. Muetzelfeldt. Position paper on declarative modelling in ecological and environmental research. Technical report, European Commission, 2004.

R. Muetzelfeldt and J. Massheder. The Simile visual modelling environment. European Journal of Agronomy, 18(3-4):345–358, 2003.

J. Nash and J. Sutcliffe. River flow forecasting through conceptual models part I – a discussion of principles. Journal of Hydrology, 10(3):282–290, 1970.

P. Papajorgji, H. W. Beck, and J. L. Braga. An architecture for developing service-oriented and component-based environmental models. Ecological Modelling, 179(1):61–76, 2004.

R. Powers. An object-oriented approach to managing model complexity. Master’s thesis, University of Bergen, 2011.

C. Pradal, S. Dufour-Kowalski, F. Boudon, C. Fournier, and C. Godin. OpenAlea: a visual programming and component-based software platform for plant modelling. Functional Plant Biology, 35(9-10):751–760, 2008.

J. M. Rahman, S. P. Seaton, and S. M. Cuddy. Making frameworks more useable: using model introspection and metadata to develop model processing tools. Environmental Modelling and Software, 19(3):275–284, 2004.

C. Rappoldt and D. van Kraalingen. The Fortran Simulation Translator FST version 2.0. Introduction and reference manual. In Quantitative approaches in systems analysis Quantitative approaches in systems analysis Quantitative approaches in system analysis, 5, pages 1–178. AB-DLO, 1996.

J. F. Reynolds and B. Acock. Modularity and genericness in plant and ecosystem models. Ecological Modelling, 94(1):7–16, 1997.

B. Richmond. STELLA: Software for bringing system dynamics to the other 98%. In International Conference of the System Dynamics Society, pages 706–718, 1985.

A. Schnepf, D. Leitner, M. Landl, G. Lobet, T. H. Mai, S. Morandage, C. Sheng, M. Zoerner, J. Vanderborght, and H. Vereecken. CRootBox: a structural-functional modelling framework for root systems. Annals of Botany, 121(5): 1033–1053, 2018.

R. Sells. Julia programming language benchmark using a flight simulation. In 2020 IEEE Aerospace Conference, pages 1–8, 2020.

R. Sequeira, R. Olson, and J. McKinion. Implementing generic, object-oriented models in biology. Ecological Modelling, 94(1):17–31, 1997.

T. D. Sharkey. What gas exchange data can tell us about photosynthesis. Plant, Cell & Environment, 39(6):1161–1163, 2016.

W. Silvert. Object-oriented ecosystem modeling. Ecological Modelling, 68(1-2): 91–118, 1993.

S. Soualiou, Z. Wang, W. Sun, P. de Reffye, B. Collins, G. Louarn, and Y. Song. Functional–structural plant models mission in advancing crop science: opportunities and prospects. Frontiers in Plant Science, 12, 2021.

D. Timlin, Y. A. Pachepsky, and B. Acock. A design for a modular, generic soil simulator to interface with plant models. Agronomy Journal, 88(2):162–169, 1996.

F. K. van Evert and G. S. Campbell. CropSyst: A collection of object-oriented simulation models of agricultural systems. Agronomy Journal, 86(2):325–331, 1994.

M. K. van Ittersum, P. A. Leffelaar, H. van Keulen, M. J. Kropff, L. Bastiaans, and J. Goudriaan. On approaches and applications of the Wageningen crop models. European Journal of Agronomy, 18(3-4):201–234, 2003.

D. van Kraalingen. The FSE system for crop simulation. In Simulation Reports CABO-TT, 23, pages 1–77. Centre for Agrobiological Research (CABO) and Department of Theoretical Production Ecology (TPE), 1991.

D. van Kraalingen. The FSE system for crop simulation, version 2.1. In Quantitative approaches in systems analysis, 1, pages 1–58. DLO Centre for Agrobiological and Soil Fertility Research (AB-DLO), 1995.

D. van Kraalingen and F. W. T. P. de Vries. The FORTRAN version of CSMP MACROS (Modules for Annual CROp Simulation). In Simulation Reports CABO-TT, 21, pages 1–145. Centre for Agrobiological Research (CABO), 1990.

D. van Kraalingen, C. Rappoldt, and H. H. van Laar. The Fortran simulation translator, a simulation language. European Journal of Agronomy, 18(3-4): 359–361, 2003.

D. W. G. van Kraalingen, M. J. R. Knapen, A. de Wit, and H. L. Boogaard. WISS a Java continuous simulation framework for agro-ecological modelling. In Environmental Software Systems. Data Science in Action, pages 242–248, 2020.

F. Villa. Integrating modelling architecture: a declarative framework for multi-paradigm, multi-scale ecological modelling. Ecological Modelling, 137(1):23–42, 2001.

J. Vos, J. B. Evers, G. H. Buck-Sorlin, B. Andrieu, M. Chelle, and P. H. B. de Visser. Functional-structural plant modelling: a new versatile tool in crop science. Journal of Experimental Botany, 61(8):2101–2115, 2010.

V. Wenzel. Semantics and syntax elements of a unique calculus for modelling of complex ecological systems. Ecological Modelling, 63(1-4):113–131, 1992.

K. Yun, J. Hsiao, M.-P. Jung, I.-T. C. Choi, D. M. Glenn, K.-M. Shim, and S.-H. Kim. Can a multi-model ensemble improve phenology predictions for climate change studies? Ecological Modelling, 362:54–64, 2017.

K. Yun, D. Timlin, and S.-H. Kim. Coupled gas-exchange model for c_4_ leaves comparing stomatal conductance models. Plants, 9(10):1358, 2020.

K. Yun, K. H. Moon, and S.-H. Kim. Cropbox for teaching: a modeling framework for teaching crop modeling and physiology. HortScience, 56(9):S34, 2021.

K. Yun, M. Shin, K. H. Moon, and S.-H. Kim. An integrative process-based model for biomass and yield estimation of hardneck garlic (Allium sativum). Frontiers in Plant Science, 2022.

X.-R. Zhou, A. Schnepf, J. Vanderborght, D. Leitner, A. Lacointe, H. Vereecken, and G. Lobet. CPlantBox, a whole-plant modelling framework for the simulation of water- and carbon-related processes. in silico Plants, 2(1):diaa001, 2020.

